# The KCNQ4-mediated M-current regulates the circadian rhythm in mesopontine cholinergic neurons

**DOI:** 10.1101/2020.09.11.293423

**Authors:** T. Bayasgalan, S. Stupniki, A. Kovács, A. Csemer, P. Szentesi, K. Pocsai, L. Dionisio, G. Spitzmaul, Pál B.

## Abstract

The M-current is a voltage gated potassium current inhibited by muscarinic activation and affected by several other G-protein coupled receptors. Its channels are formed by KCNQ subunits, from which KCNQ4 is restricted to certain brainstem structures. We sought evidence for the function of the M-current in the pedunculopontine nucleus (PPN) and the contribution of KCNQ4 subunits to the M-current and aimed to find its functional significance in the PPN. We found that cholinergic inputs of the PPN can effectively inhibit M-current. This current is capable of synchronizing neighboring neurons and inhibition of the M-current decreases neuronal synchronization. We showed that only a subpopulation of cholinergic neurons has KCNQ4-dependent M-current. The KCNQ4 subunit expression potentially regulates the presence of other KCNQ subunits. Deletion of KCNQ4 leads to alterations in adaptation of activity to light-darkness cycles, thus representing the potential role of KCNQ4 in regulation of sleep-wakefulness cycles. The presence of this protein restricted to certain brainstem nuclei raises the possibility that it might be a potential target for selective therapeutic interventions affecting the reticular activating system.

## Introduction

The M-current is a voltage-gated potassium current, which is under the regulation of several neuromodulatory actions and receptors. Probably the best-known modulatory action on these channels is the inhibition by muscarinic acetylcholine receptors (Brown and Adams, 1980; Hernandez et al, 2008; Marrion, 1997). The neuronal M-current sets resting membrane potential, regulates excitability and shapes action potential firing (Brown and Passmore, 2009; Delmas and Brown, 2005). In presynaptic location, it controls synaptic vesicle release (Huang and Trussell, 2011).

Ion channels responsible for M-current are composed of KCNQ (Kv7) subunits, which can form homo- or heterotetrameric channels. KCNQ2-5 subunits are found in the central nervous system (CNS), from which KCNQ2, 3 and 5 are abundantly located in several brain areas, whereas KCNQ4 is restricted to certain nuclei of the brainstem (Brown and Passmore, 2009; Delmas and Brown, 2005, Kharkovets 2000). These comprise of auditory brainstem nuclei as the cochlear nuclei, nuclei of the lateral lemniscus and the inferior colliculus; the principal and spinal trigeminal nuclei; and members of the reticular activating system (RAS) as raphe nuclei and the ventral tegmental area (Kharkovets et al, 2000; Hansen et al 2008; Koyama and Appel, 2006).

Mutations in the *KCNQ4* gene leads to an autosomal progressive nonsyndromic hearing loss due to the degeneration of outer, and in a lesser extent, inner hair cells of the cochlea, known as DFNA2 (De Leenheer et al, 2002; Nie et al, 2008). The disease has been reproduced in mouse models that either expressed a human KCNQ4-mutation or lacked KCNQ4 channel expression (Kharkovets et al., 2006, Carignano et al 2019). The mutation alters somatosensory functions as well due to the lack of expression on skin somatosensory receptors and dorsal root ganglia either in human and mouse (DRG, Heidenreich et al, 2011).

It can be well seen that KCNQ4-positive brainstem areas overlap with cholinoceptive areas (Woolf and Butcher, 2011; Kharkovets, 2000) and that the M-current is affected by cholinergic neuromodulation (Brown and Passmore, 2009; Delmas and Brown, 2005). It has been also recently shown that the M-current formed by KCNQ4 subunits contribute to neuromodulatory autoregulation (Su et al, 2019). Therefore, we sought evidence for the hypothesis that M-current formed by KCNQ4 subunits is present on mesopontine cholinergic neurons.

One of the main mesopontine cholinergic areas is the pedunculopontine nucleus (PPN) which is not only the source of cholinergic fibers but receives cholinergic inputs from the neighboring laterodorsal tegmental nucleus (LDT) and the contralateral PPN. Besides cholinergic inputs from neighboring structures, local cholinergic collaterals exist in the nucleus (Mena-Segovia et al, 2008; Honda and Semba, 1995). The PPN has cholinergic and non-cholinergic neurons (GABAergic, glutamatergic neurons), which show different activity pattern during modulation of global brain states. The PPN has distinct neuronal activity pattern in slow wave sleep (SWS), paradoxical sleep (PS) and wakefulness (W). Briefly, PPN units are synchronized during SWS but the level of synchronization is reduced during PS and W (Petzold et al, 2015; Mena-Segovia and Bolam, 2017). As local neuronal networks of the PPN have not been mapped well, mechanisms of this synchronization have not been presented yet in detail.

We previously showed that almost all PPN cholinergic neurons possess M-current, whereas GABAergic neurons lack it. The M-current of the cholinergic neurons is responsible for the afterhyperpolarization current and spike frequency adaptation, and is capable of modulating high threshold membrane potential oscillations (Bordás et al, 2015). However, there are still some pending questions regarding the M-current of the PPN.

We aimed to confirm that the M-current is the hallmark of PPN cholinergic neurons, and demonstrate that KCNQ4, which is specific to the RAS, is present on these neurons. Besides, we also evaluated the physiological significance of the M-current in the PPN.

We showed that the M-current exists on the soma of cholinergic neurons and not on glutamatergic ones, and activation of cholinergic inputs of the PPN can inhibit its M-current. The M-current contributes to synchronization of neighboring neurons. We demonstrated the presence of KCNQ4 on a subgroup of cholinergic neurons by using several methods. We also showed that KCNQ4-knock-out (KO) mice displayed alterations in activity cycles. Our findings add new roles for the CNS-expressed KCNQ4 channel such us determining the molecular composition of the KCNQ channels in the RAS and thus, its electrophysiological properties that, in turn, modulates the activity cycles. As the expression of this subunit is restricted to certain brainstem nuclei, subunit-specific modulators can potentially act as medication for disturbances in sleep-wakefulness cycles.

## Results

### 1. Insights for M-current localization in neurons from PPN

#### 1.1. Neuronal population analysis

We previously hypothesized that the M-current is exclusively present on cholinergic neurons in the PPN, based on data obtained from genetically labelled cholinergic (ChAT+) and GABAergic (GAD65+), but not glutamatergic neurons (Bordás et al, 2015). First, we aimed to confirm it by analyzing M-current in genetically labelled PPN glutamatergic neurons (Vglut2+; Fig. 1A). As expected, the M-current was present in almost all ChAT+ PPN neurons and absent in the GAD65+ ones (Fig. 1B-D). In glutamatergic neurons, 6.8% of them (n=32) presented the M-current. One of these neurons proved to be ChAT-positive, possibly belonging to the population of glutamatergic-cholinergic neurons (Wang and Morales; 2009; Baksa et al, 2019). We concluded that the majority of non-cholinergic neurons lack the M-current, but very few exceptions may exist for the glutamatergic neuron population (Fig. 1E).

**Fig.1.**
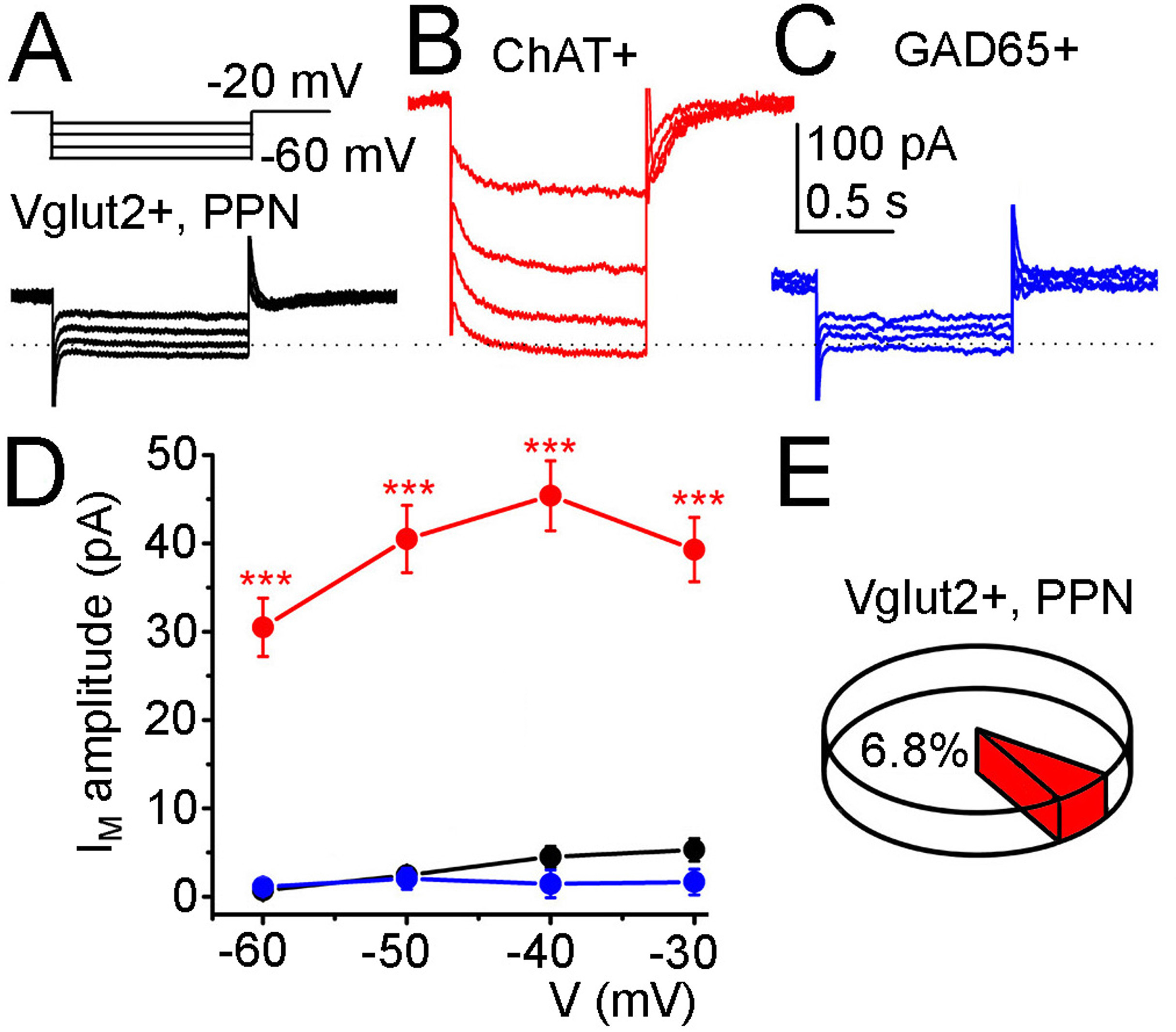
Only cholinergic and not glutamatergic and GABAergic PPN neurons possess M-current. A-C. Current traces recorded from a glutamatergic (A, inset above the current trace: voltage protocol detailed in Methods), cholinergic (B) and a GABAergic neuron (C). D. Statistical comparison of the M-current recorded from a cholinergic (red), glutamatergic (black) and a GABAergic neuron (blue) in the function of the repolarizing voltage steps (*** shows significant difference from Vglut2+ at p<0.001; dotted line: 0 pA). E. Percentage of glutamatergic neurons possessing M-current.

#### 1.2. Subcellular localization of the M-current

To clarify the location of the M-current in the cholinergic neurons, we performed recordings from the proximal dendrite (Fig. 2A, C) and the soma (Fig. 2B, D) of the same neuron one after the other in 17 cases (Fig. 2). The holding current at −20 mV, which is supported in part by open KCNQ channels, was significantly lower for the proximal dendrites than for the soma (1.17 ± 0.15 vs 1.67 ± 0.23 pA/pF current density, for the proximal dendrite and the soma, respectively; p=0.035). Then we determined M-currents from dendrites and soma by measuring the relaxation currents obtained at each voltage step (Fig. 2E, F). M-currents for proximal dendrites were significantly lower than somatic ones at all voltages (0.158 ± 0.03 vs. 0.308 ± 0.06 pA/pF, p = 0.01 at −60 mV; 0.253 ± 0.04 vs 0.409 ±0.07 pA/pF, p = 0.03 at −50 mV; 0.302 ± 0.047 vs 0.484 ± 0.09 pA/pF, p = 0.03 at −40 mV; 0.267 ± 0.04 vs. 0.418 ± 0.084 pA/pF, p = 0.044 at −30 mV; for the proximal dendrite and the soma, respectively; Fig.2G). Taken together, KCNQ channel location seems to be higher in the soma than the proximal dendrites, or it could reflect different molecular composition. All of the further patch clamp recordings were performed from the soma.

**Fig.2.**
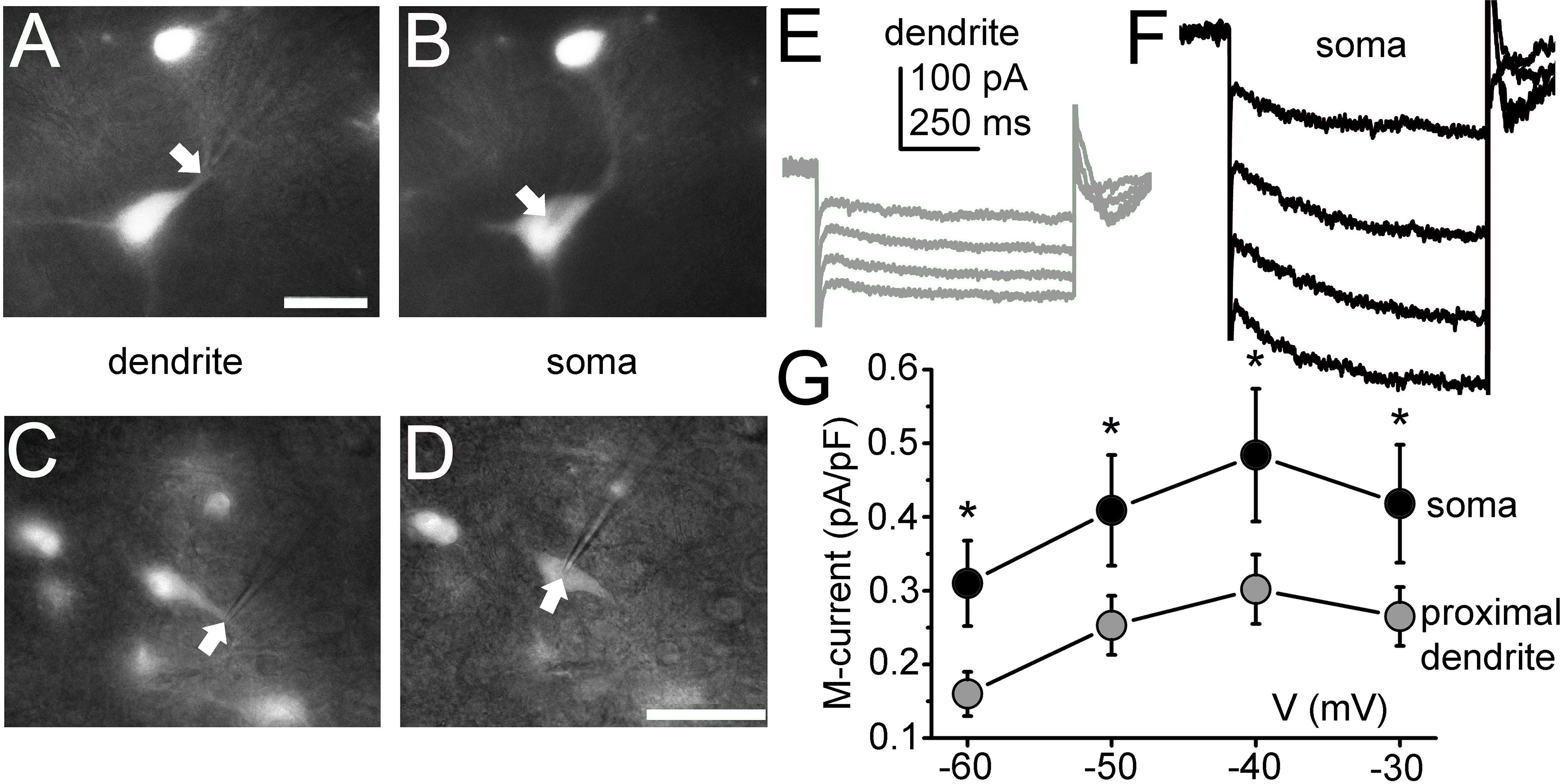
The recorded M-current has a significantly greater amplitude on neuronal somata than on dendrites. A-D. Two examples for a dendritic (A, C) and somatic (B, D) patch clamp recordings on ChAT-tdTomato samples. Photomicrographs of panels A-B and C-D were taken on the same neurons, respectively. E-F. M-current recorded from the dendrite (E, grey) and the soma (F, black) from the same neuron. G. Statistical summary of the M-current density recorded from the soma (black) and dendrite (grey) in the function of the repolarizing voltage steps (* shows significant difference from proximal dendrite at p<0.05.) Scale bars: 50 μm.

#### 1.3. M-current modulation in cholinergic neurons

In the next set of experiments, we aimed to test whether a near-physiological activation of a cholinergic input of the PPN neurons like the laterodorsal tegmental nucleus (LDT), can effectively inhibit the M-current on PPN cholinergic neurons. In this experiment, LDT of ChAT-ChR2 mice was optogenetically stimulated with 1 Hz frequency in a 500-μm-thick coronal brainstem block while the M-current of PPN cholinergic neurons was recorded (Fig.3A-C). The holding current decreased to 60.45 ± 6.49% of control (p= 0.008; Fig.3C-D). The relaxation current also decreased at all voltages tested. The magnitude of the change was in between 51-73% of control (56.11 ± 12.75% at −60 mV, 51.1 ± 15.27% at −50 mV, 62.57 ± 12.53% at −40 mV, 72.7 ± 24.17% at −30 mV; Fig.3E). 5 minutes after the illumination, the M-current was partially regenerated (88.6 ± 21.6 pA holding current, p = 0.093; 7.46 ± 5.15 pA M-current amplitude at −60 mV, p = 0.089; 24.63 ± 5.27 pA at −50 pA, p = 0.05; 35.07 ± 10.78 pA at −40 pA, p = 0.07; 33.57 ± 4.39 pA at −30 pA, p = 0.04). We concluded that optogenetic activation of a single cholinergic input of the PPN can effectively inhibit the M-current.

**Fig.3.**
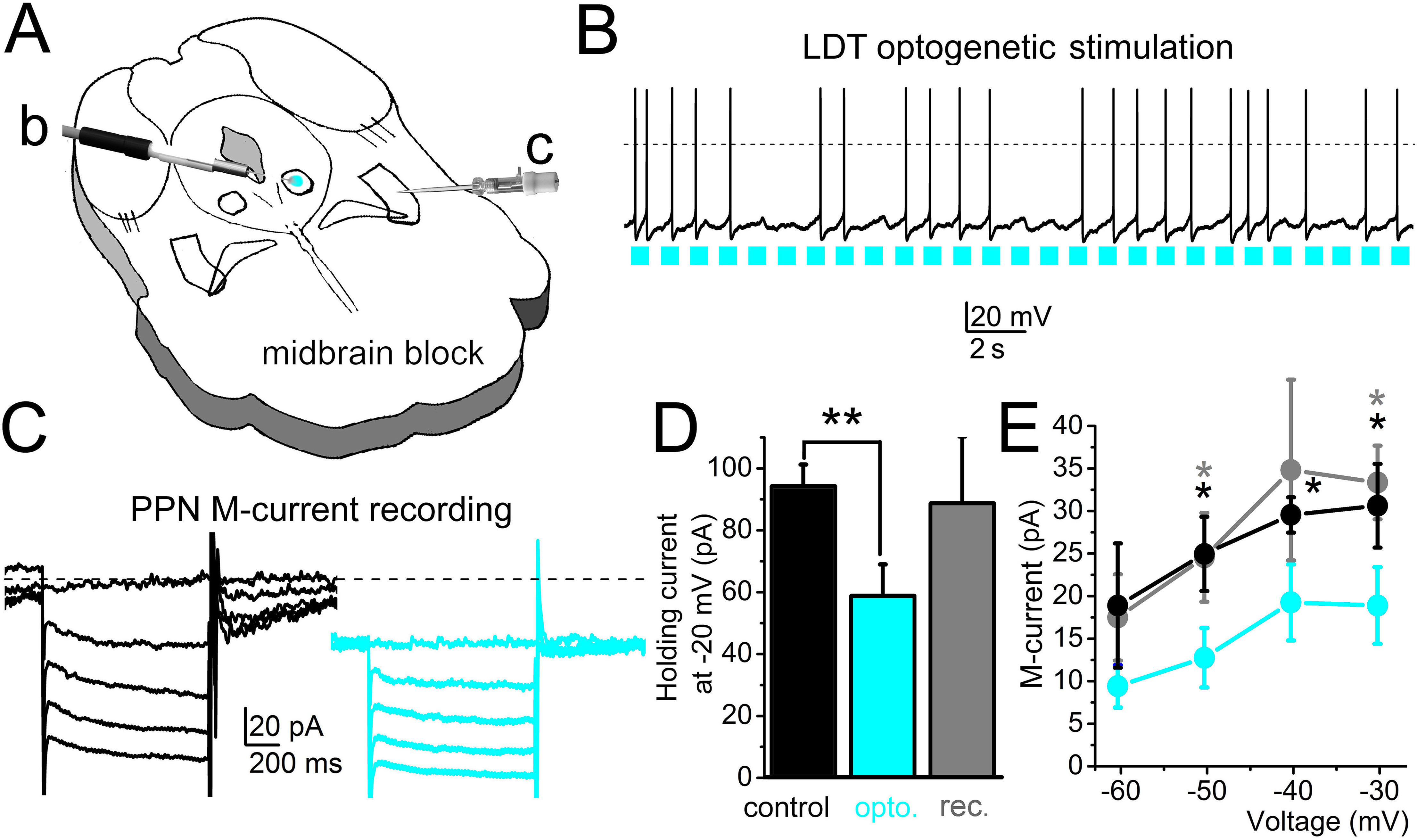
Stimulation of a cholinergic PPN input can effectively reduce the PPN M-current amplitude. A. Schematic drawing of the experimental arrangement. The laterodorsal tegmental nucleus (LDT) of the midbrain block prepared from a ChAT-ChR2 mouse was optogenetically stimulated (blue spot; b; see panel B) in parallel with patch clamp recording from the PPN (c; see panel C). B. Optogenetic stimulation of a cholinergic neuron in the LDT with 1 Hz pulsatile blue light (blue squares) and the consequential action potential firing of the activated neuron. C. The M-current recorded from a PPN cholinergic neuron before (black) and during (blue) optogenetic stimulation of the LDT. D. Statistical comparison of the holding currents at −20 mV voltage recorded before (black), during optogenetic stimulation (blue) and after recovery (gray; **: p<0.01). E. Statistical comparison of the repolarizing current steps recorded before (black), during optogenetic stimulation (blue) and after recovery (gray; *: p<0.05.)

Next, we tested whether blockade of the M-current can alter synchronization in firing activity of neighboring PPN neurons. Namely, the spike frequency adaptation (SFA) was measured by using the adaptation index. Inhibition of the M-current by XE991 reduced the adaptation index from 0.353 ± 0.026 to 0.277 ± 0.028 (p = 0.0375). Two synaptically non-coupled neurons were depolarized simultaneously with 100 pA, action potential trains were recorded and time differences between the closest action potentials were assessed (Fig.4A-G). Values of delays between the action potentials of the neighboring neurons showed an altered distribution when M-channel is absent (Fig.4H). In control, more than 50% of the delay values are shorter than 40 ms. When XE991 is present, delay values are more uniformly distributed for the first 100 ms (Fig. 4H). The level of synchronicity in the presence of the M-current blocker decreased significantly (Fig. 4I).

**Fig.4.**
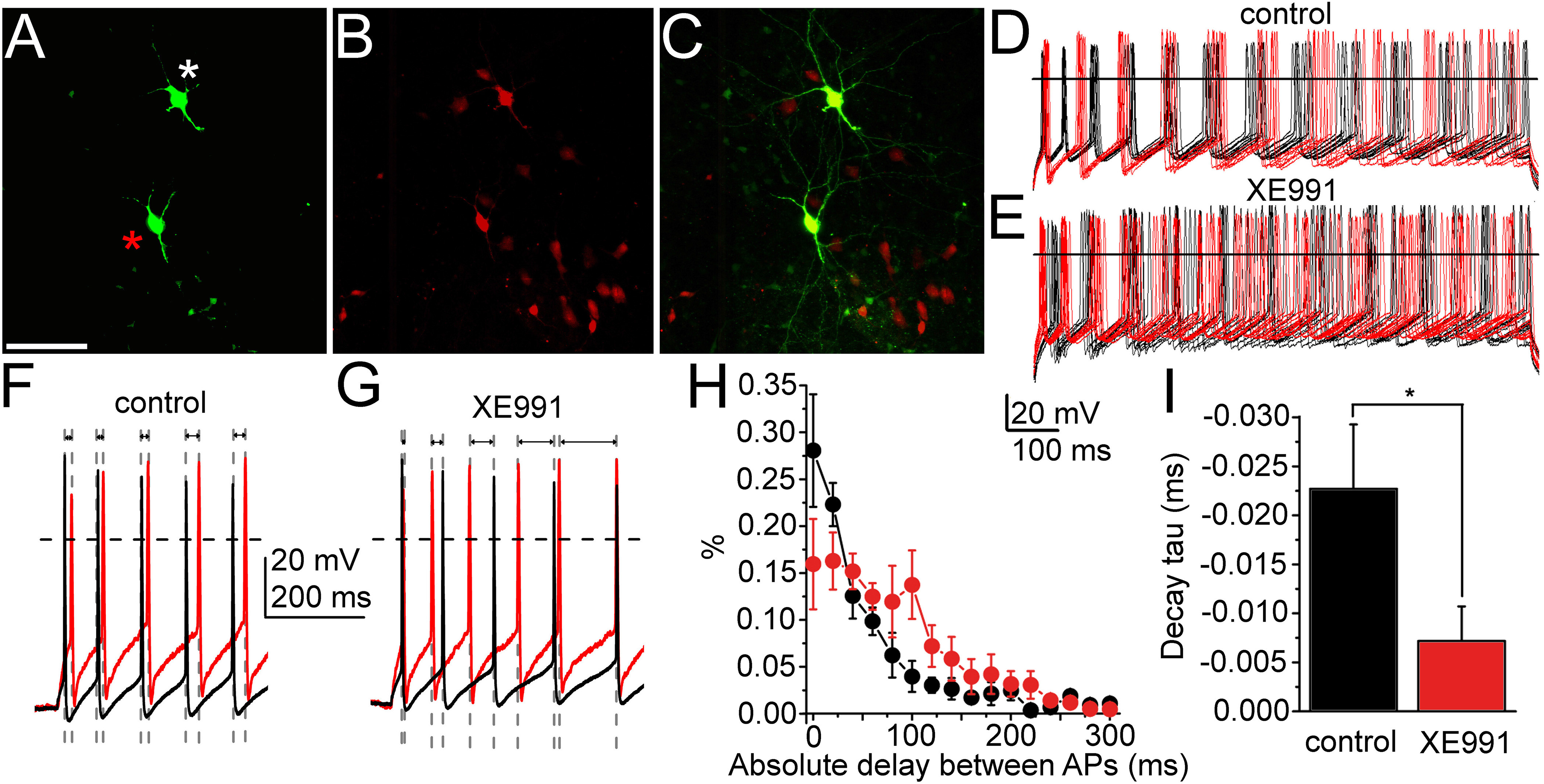
The M-current of PPN cholinergic neurons can control synchronization of the neighboring neurons. A-C. Two cholinergic neurons labelled in the PPN during recordings. A. Biocytin labelling (scale bar: 100 μm). B. ChAT-dependent tdTomato expression. C. Merged image. D-E. Action potential series elicited by 100 pA depolarizing stimuli from the two neighboring neurons under control conditions (D, red traces: recordings from the neuron labelled with red asterisk; black traces: recorded from the neuron labelled with white asterisk) and in the presence of the M-current inhibitor XE991 (E). F-G. Representative pairs of traces at high time resolution under control conditions (F) and with XE991 (G) representing changes of delay intervals between spikes (indicated with gray dashed lines and arrows). H. Average histograms of absolute delay between action potentials recorded from the neighboring neurons with 20 ms bins (average ± SEM; black: control, red: with XE991). I. Decay tau values obtained with the fit of the individual histograms of absolute delays under control conditions (black column) and with XE991 (red column).

With these results, we confirmed our previous findings that the M-current exists on cholinergic and is absent from non-cholinergic neurons of the PPN. We also showed that mostly somatic M-current can be effectively inhibited by the physiological cholinergic inputs of the nucleus. Blockade of M-current causes loss of synchronization between neighboring neurons.

### 2. Contribution of KCNQ4 channel subunit to the M-current in PPN neurons

#### 2.1. Expression of KCNQ4 channel subunit in PPN

It was shown that KCNQ4 channel is present in some nuclei of the RAS so we hypothesized that the PPN, as a member of the RAS and subject of cholinergic neuromodulation via muscarinic activation and M-current inhibition, might also possess this subunit. To prove this, we analyzed KCNQ4 expression in mice by qPCR and its localization using immunofluorescence, and the consequences of KCNQ4 lacking on the other KCNQ subunits. We evaluated the expression pattern of neuronal KCNQ subunits at mRNA level obtained from the PPN in KCNQ4-KO and wild type (WT) animals (Fig.5A-C). We observed expression of *Kcnq2* to *Kcnq5* subunits in the PPN with *Kcnq2* showing the highest mRNA expression in WT animals. In KO mice, the expression profile changed drastically: *Kcnq2* decreased to an expression level similar to *Kcnq3* and the mRNA of *Kcnq5* disappeared (Fig. 5B). To relatively quantify these changes in mRNA expression, we calculated the fold change (RQ) of each subunit comparing WT with KO animals. We observed more than a 35-fold decrease for *Kcnq2* subunit mRNA expression in KO animals (p=0.032, One-way ANOVA, *post hoc* Tukey test, for both genotypes; Fig. 5C). On the contrary, the expression level of *Kcnq3* in KO was about 7-fold higher than in WT animals (p<0.010, Student’s t-test; Fig. 5C).

**Fig.5.**
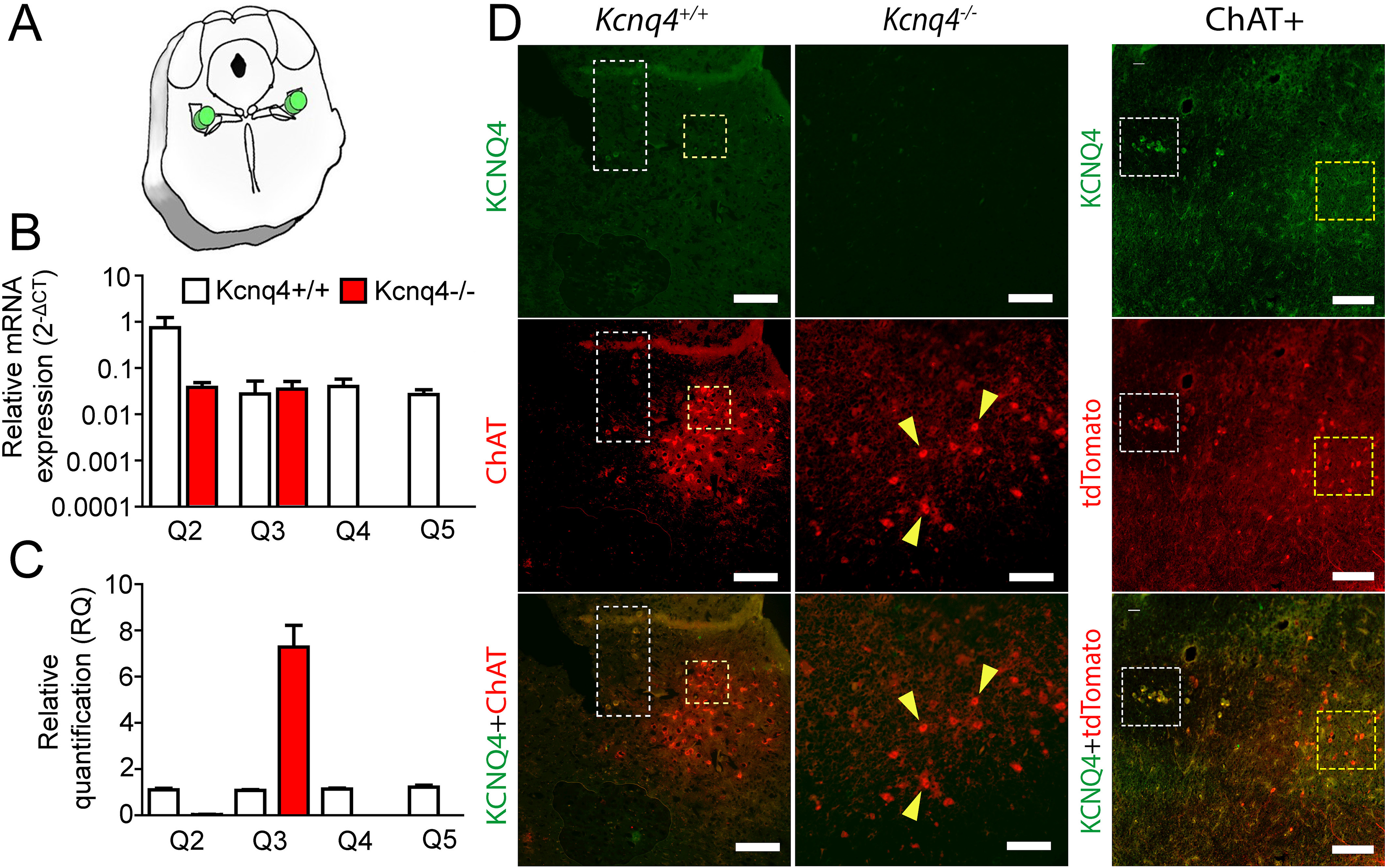
KCNQ4 lacking alters the mRNA expression of the other KCNQ subunits. KCNQ4 is located only in a subpopulation of PPN cholinergic neurons. A. Location of cylinder-shaped tissue biopsy (green symbol) from 1-mm-thick coronal midbrain slices (−5.2 - −4.2 mm from bregma). B. KCNQ2-5 subunits mRNA expression profile of WT (*Kcnq4*+/+; white) and KCNQ4-KO (*Kcnq45* −/−; red) mice PPN. C. Relative quantification (RQ) of mRNA expression for each subunit from WT (white) and KO (red) mice. The fold change of each subunit was calculated using 2^-ΔΔCT^ (data represented as mean ± SD; ***, p < 0.001). D. KCNQ4 immunofluorescence on coronal brain sections of WT (*Kcnq4*+/+; left panel) and ChAT-tdTomato (ChAT+; right panel) mice. Both models revealed that only a subpopulation of PPN cholinergic neurons located on the external limits possess KCNQ4 (white dashes square), and the rest of them were negative for the subunit (yellow dashed square). KCNQ4 could not be detected on PPN cholinergic neurons (yellow arrows) in KO (*Kcnq4*−/−) animals, that confirmed the specificity of the antibody. (Upper panel: KCNQ4 (green), Middle panel: ChAT immunolabelling or tdTomato expression under ChAT promoter (red). Bottom panel: Merged image. Scale bar: 50 μm, 20 μm and 50 μm, for each column).

These results show that the absence of KCNQ4 in PPN alters the expression pattern of KCNQ channel subunits and probably may change the properties of the M-current in KO animals.

In order to check the presence of KCNQ4 as well as for determining its localization, KCNQ4 immunofluorescence was performed on WT mice using anti-ChAT antibody to identify cholinergic neurons and in a transgenic mouse with genetically labelled cholinergic neurons. In both cases, we found KCNQ4 in a subset of cholinergic neurons (Fig. 5D). In most of them, KCNQ4 signal was present in the whole cytoplasm and in minor cases seems to be restricted to the somatodendritic surface membrane. KCNQ4 is rather present in caudal than rostral locations. Even in caudal location, not the whole population exhibited KCNQ4-specific labeling, only laterally located subgroups (Fig. 5D). KCNQ4 signal is absent in PPN section of KCNQ4-KO animals, but the number of cholinergic neurons is rather similar to WT (Fig. 5D). After cell count, we determined that the proportion of KCNQ4-positive PPN cholinergic neurons was 9.0 ± 4.8 % (n = 4) in WT and ChAT-tdTomato mice.

#### 2.2. M-current properties in KCNQ4 KO mice

To provide support for the previous experiments, we tested whether PPN cholinergic neurons of KCNQ4 KO animals possess M-current. In this experiment, we patched PPN neurons from samples of KCNQ4 KO and WT animals. Neurons were labelled with biocytin and the cholinergic nature of neurons was checked with *post hoc* ChAT immunohistochemistry. The holding current at −20 mV was much lower in KO animals than WT. Average values were 175.97 ± 27.67 pA for the WT and 66.95 ± 19.8 pA for the KO animals (p = 0.006) (Fig. 6A). In addition, we determined that the M-current was absent or highly decreased (< 10 pA) in 62.5% of the KO cases while WT animals only exhibited its absence in 7.7% of the cases (n = 13 for WT, and n = 8 for KO; Fig.6A, B). The average relaxation current at −40 mV was 42.57 ± 7.95 pA in the WT and 14.85 ± 6.12 pA in the KO animals (p = 0.013; Fig. 6C).

**Fig.6.**
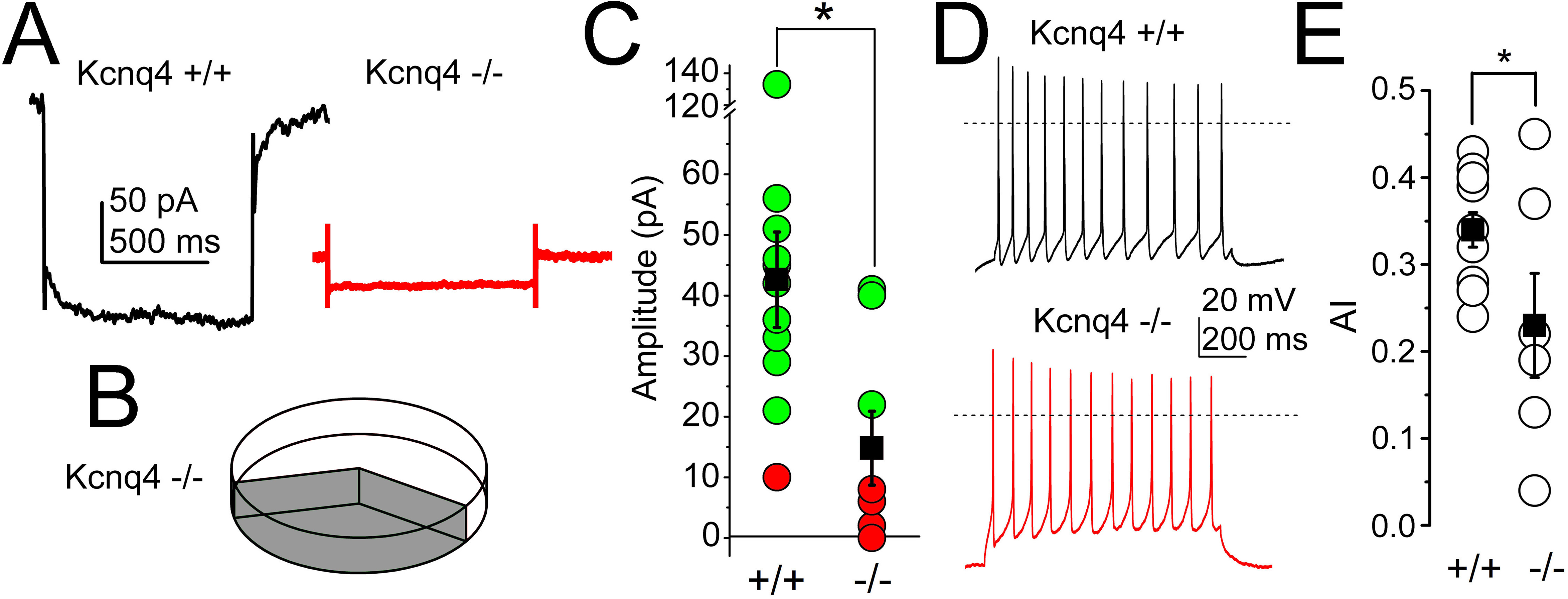
More than 60% of the cholinergic neurons lack M-current in KCNQ4-KO samples. A. Representative current traces from a WT (*Kcnq4*+/+, black) and KO (*Kcnq4*−/−, red) samples. B. Percentage of cholinergic neurons from KO samples lacking (hollow) and possessing (gray) M-current. C. Statistical comparison of relaxation current at −40 mV from WT (+/+) and KO (−/−) samples (green: M-current exists, red: no M-current detected; black squares: average ± SEM). D. Representative voltage traces obtained from WT (*Kcnq4*+/+, black) and KO (*Kcnq4*−/−, red) samples. Dotted lines indicate 0 mV. E. Statistical comparison of adaptation index (AI) from WT (+/+) and KO samples (−/−; hollow circles: individual data, black squares: average ± SEM; * shows significant difference at p<0.05.)

As the M-current exerts an important action on the SFA, being capable of determining synchronization of the neighboring neurons, we analyzed whether the absence of KCNQ4-mediated M-current can affect SFA. We elicited trains of action potentials by depolarizing square current injections and then we calculated the adaptation index (AI) of the trains. In most cases, KCNQ4 KO mice exhibited a slight higher amount of action potentials generated due to current injection, when compared to WT animals (Fig.6D). The AI of cholinergic neurons from KCNQ4 KO animals were significantly lower than WT (p = 0.039; n = 6 for KO and n = 9 for WT; Fig. 6E). However, in agreement with the experiments shown above, 66.67% of the KO neurons exhibited lower AI.

In summary, these experiments showed that KCNQ4 is present in a minor proportion of neurons in PPN (together with KCNQ2, KCNQ3 and KCNQ5), but its absence alters the expression of the other subunits and the M-current properties as well as its functional role.

#### 2.3. Presence of functional KCNQ4 subunits in PPN

In order to discriminate the function of KCNQ4 in PPN-cholinergic neurons under WT conditions, we dissected its contribution by using different subunit-specific KCNQ channel openers. For control, we performed holding current analysis using 20 μM retigabine, a non-specific KCNQ channel opener. Retigabine elicited an outward shift of the holding current at −20 mV on 100% of the neurons (n = 11; Fig.7A, D). The holding current at −20 mV increased from 95.25 ± 11.96 pA to 140.23 ± 12.24 pA (p= 0.009; Fig. 7A, E).

**Fig.7.**
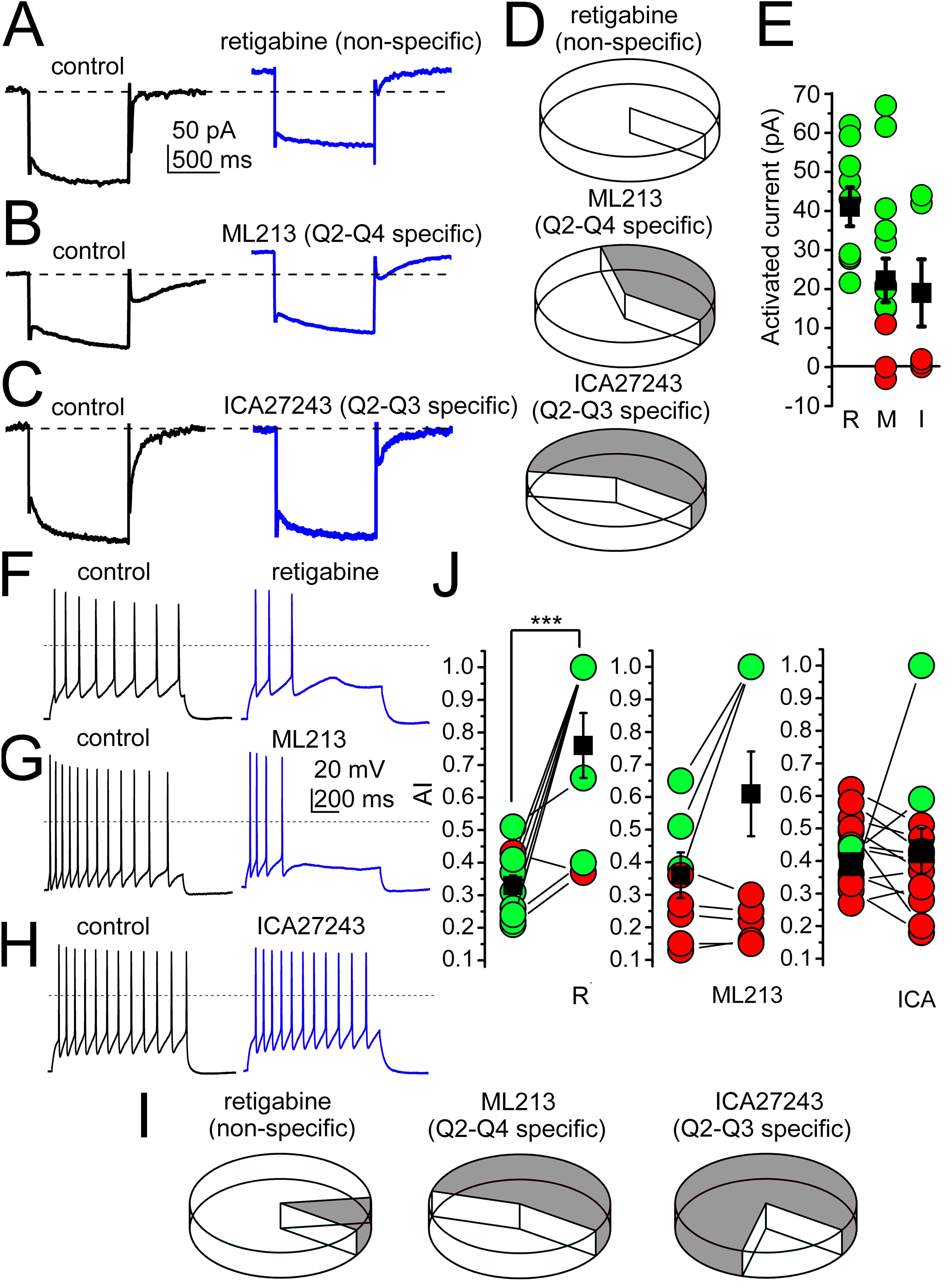
The presence of the KCNQ4 subunit can be detected on a subpopulation of PPN cholinergic neurons. A. Actions of the non-specific KCNQ opener retigabine on the M-current (black: control trace, blue: trace with retigabine). B. Representative M-current traces under control conditions (black) and with the KCNQ2- and KCNQ4-specific opener ML213 (blue). C. Representative M-current traces under control conditions (black) and with the KCNQ2- and KCNQ3-specific opener ICA27243 (blue). Dashed lines indicate the holding currents in control. D. Percentage of neurons with M-current activation by the drug presented above (hollow: M-current activation, gray: no-M-current activation). E. Statistical comparisons of currents activated by retigabine (R), ML213 (M) and ICA27243 (I) at −20 mV holding potential (green: outward currents greater than 10 pA, red: currents less than 10 pA, black squares: average ± SEM). F-H. Representative voltage traces obtained with 100 pA depolarizing current injections under control conditions (black) and with non-selective (F, retigabine), KCNQ2- and KCNQ4-selective (G, ML213) and KCNQ2- and KCNQ3-selective (H, ICA27243) openers. Dotted lines indicate 0 mV. I. Percentage of PPN cholinergic neurons increasing adaptation index (AI) to various M-current openers (hollow) and lacking response (gray). J. Statistical comparison of changes in AI by openers with different selectivity (green: increase, red: no change or decrease in individual data; black squares: average ± SEM; *** show significant difference at p<0.001).

Next, we tested the KCNQ2- and KCNQ4-specific M-current opener ML213 (20 μM). This generated a shift in the holding current in 69.2% of the cholinergic neurons tested and the rest were insensitive to it (Fig. 7B, D). The overall average increase of the outward current at −20 mV holding potential was 25.61 ± 5.94 pA (from 143.99 ± 28 pA to 169.61 ± 29.66 pA; n =13; Fig. 7 E).

Then, we administered the KCNQ2- and KCNQ3-specific opener ICA27243 (20 μM). In the majority of the cases, this opener did not change holding current (Fig. 7C, D). Signs of KCNQ channels activation were seen on 42.86% of the neurons (n = 7; Fig. 7E). It elicited an averaged additional increase in the outward current of 19 ± 8.61 pA at −20 mV holding potential (from 165.04 ± 23.23 pA to 174.96 ± 25.78 pA; Fig 7E; Table.1).

**Table.1.**
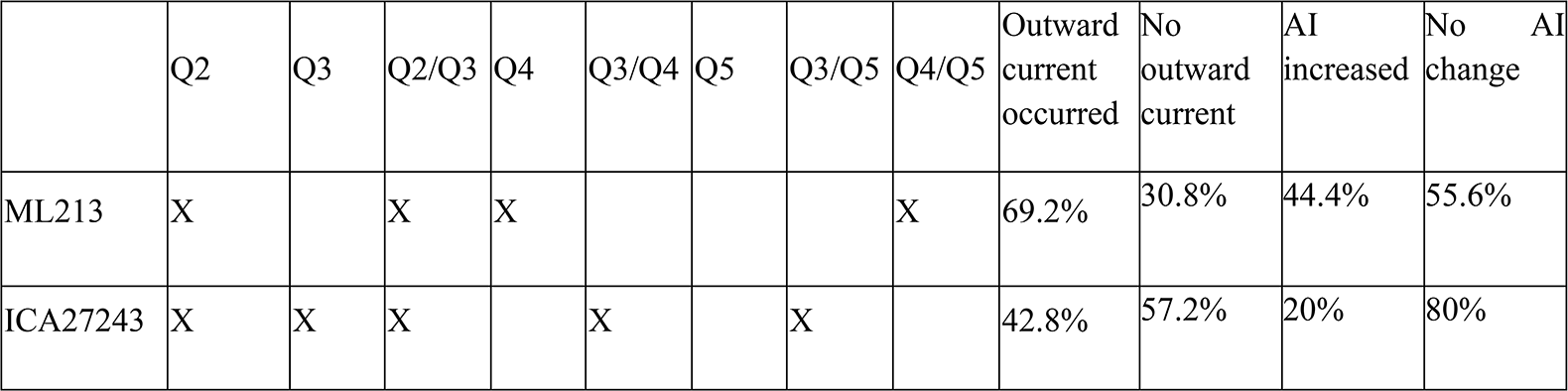
Actions of M-current openers on KCNQ subunits and on potential heteromers and percentages of cholinergic neurons responding to the specific openers in different experiments.

In the next series of experiments, we evaluated the effect of the openers on the AI. With retigabine, we observed an increase of AI in the vast majority (88.9%) of the cases, from 0.33 ± 0.035 to 0.755 ± 0.1 (p = 0.0005; n = 9; Fig.7F, I, J), indicating that the M-current participates in the SFA of cholinergic neurons, although other channels also contribute. ML213 caused a weaker action as it increased the AI only in 44.4% of the cases (from 0.36 ± 0.06 to 0.61 ± 0.13; n.s.; n = 10; Fig.7G, I, J). With ICA27243, the effect on AI was even weaker: it increased only in 20% of all cases (from 0.39 ± 0.02 to 0.43 ± 0.07; n.s.; n = 10; Fig.7H-J; Table 1).

Taken together, we can state that not all PPN cholinergic neurons possess KCNQ4-mediated M-current. Based on percentages of activated neurons elicited with all openers, and their selectivity, we can assume that only a subpopulation of PPN cholinergic neurons possess functional KCNQ4 subunits (Table 1). This population might be approximately 25% of the whole cholinergic PPN neuronal population.

#### 2.4. Functional role of KCNQ4 in the RAS

PPN is a member of the RAS that modulates the circadian rhythm. We demonstrated the presence of KCNQ4 in this nucleus and its functional role in neuronal properties. Then, we investigated if this subunit contributes to the regulation of the circadian rhythm by the PPN by testing voluntary wheel running in young adult KCNQ4-KO and WT mice (n = 13 and 15, respectively). Mice were kept in a room with 6 hours illumination followed by 6 hours darkness (LD condition). After 7 days of adaptation and 5 days of recording, the illumination was turned off for the next 5 days (DD condition). The period time and distance were determined under both LD and DD conditions by using the Lomb-Scargle periodogram (Fig. 8A, B).

**Fig.8.**
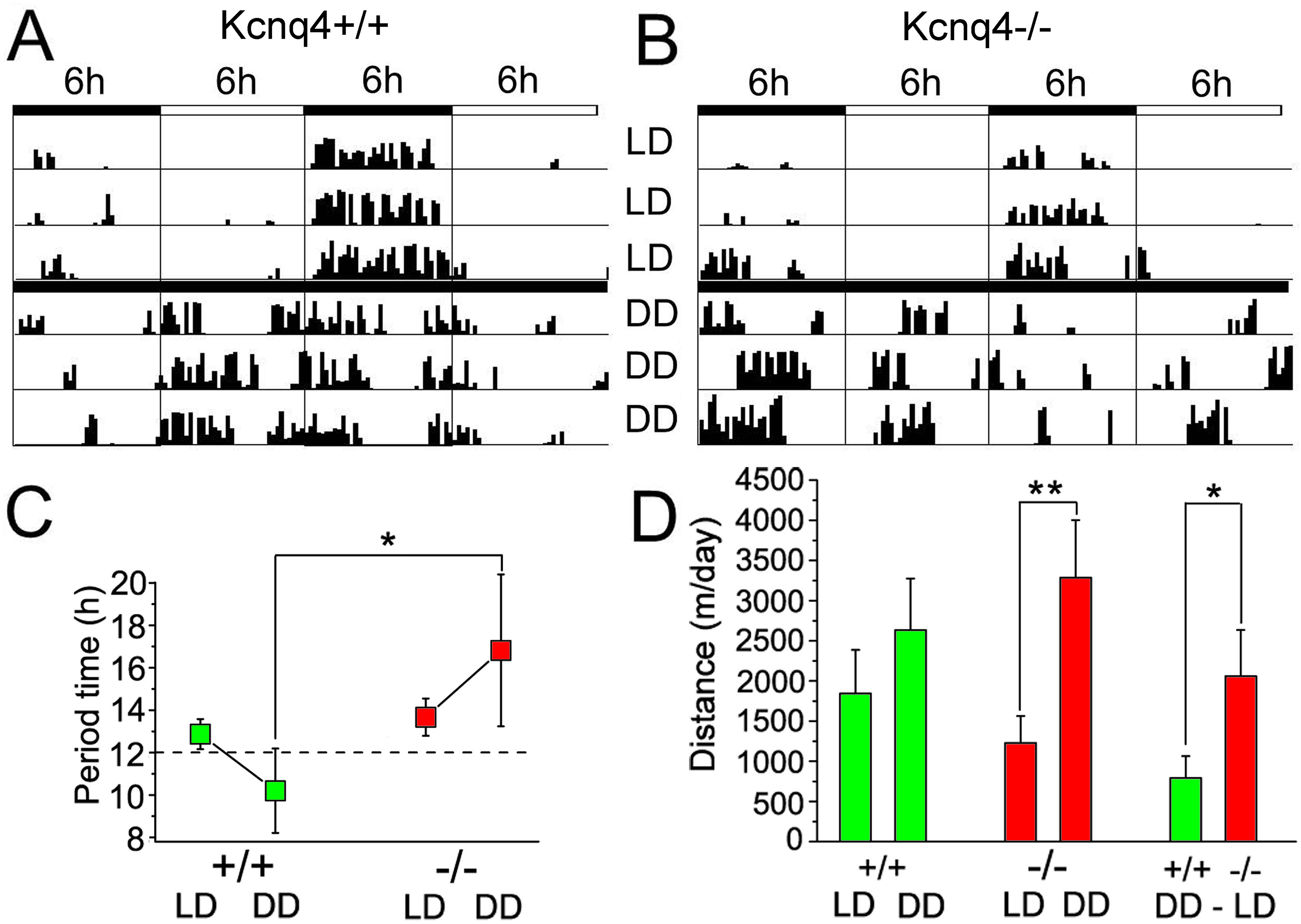
KCNQ4-KO mice show alterations of activity cycle adaptation to changes in light-darkness conditions with activity wheel test. A. Activity cycles recorded with a WT mouse (Kcnq4+/+) with 6 h alterations of light-darkness cycles (LD) and in complete darkness (DD). Black vertical bars represent distances ran in 10 minutes bins. B. Activity cycles with the same arrangement as A for KCNQ4-KO mouse (Kcnq4−/−). C. Statistical comparison of alterations in period length under DD conditions with WT (+/+) and KO (−/−) mice. D. Comparison of distances ran in 3 days by WT (+/+) and KO (−/−) under LD and DD conditions. DD-LD represents the difference in distances ran by each mice genotype under the two different conditions (*, and ** show significant difference at p<0.05, and p<0.01, respectively).

The period time under LD conditions showed no difference between genotypes. Under DD condition, the period time became significantly longer in KO mice against WT (p = 0.05; Fig.8C). For each genotype, we did not observe significant differences between LD and DD conditions.

Next we studied the distance ran by both mice genotypes in LD and DD conditions (Fig. 8D). In WT animals, we observed a tendency to increase the distance traveled in DD conditions, although it was not statistically significant. Contrary to WT, KO mice showed a ~2-fold increase in the distance traveled in DD compared to LD condition. (p = 0.009; Fig.8D). We did not find significant difference between distance traveled between genotypes neither for LD nor for DD conditions. However, the difference between the distance traveled in DD and LD conditions for each genotype was significantly different: in WT animals, the increase was 790.65 ± 274.02 m/day, whereas in KO was 2059.1 ± 575.99 m/day (p = 0.027; Fig. 8D).

Taken together, lack of KCNQ4 seems to have mild but detectable impact on adaptation to changes in light/dark cycles, as well as on movement regulation related to activity cycles.

## Discussion

Our findings assign a functional role for KCNQ4 channel expressed in the RAS and provide evidence for its expression and modulation of the M-current in the PPN.

We demonstrated that the M-current is almost exclusively present on cholinergic and almost fully absent on non-cholinergic neurons of the PPN. We also showed that the largely somatic M-current is effectively inhibited by the cholinergic inputs of the PPN. The inhibition of the M-current decreases SFA, which decreases spontaneous synchronization of the neighboring neurons. The M-current of the PPN cholinergic neurons is determined by the KCNQ4 subunit in ~a quarter of the whole population but we found that is very important for the expression of the other KCNQ-channel subunits. Deletion of KCNQ4 caused disturbances in adaptation to altered environmental light-darkness cycles.

It has been well described that the M-current is present in nuclei of the RAS, as the ventral tegmental area (VTA) and in the raphe nuclei (Drion et al, 2010; Zhao et al, 2017; Su et al 2019). We previously demonstrated that the M-current is found on PPN cholinergic neurons and controls SFA as well as other excitability-related parameters (Bordás et al, 2015). Although we provided description of the M-current of PPN neurons, several questions remained open. First, we hypothesized that the presence of M-current is a functional marker of the cholinergic neurons. Our hypothesis was based on experiments using genetically identified cholinergic and GABAergic neurons. In the present project, we added data from genetically identified glutamatergic neurons and confirmed our previous statement. Another important finding of the previous work was that the SFA is critically affected by the M-current. Here ‒in accordance with modelling and data from other brain areas (Leao et al, 2009; Roach et al, 2015; 2016)-we showed that SFA helps synchronization of neuronal populations. Blockade of the M-current decreased SFA, which decreased synchronization between two neighboring neurons. Although stimulation of a single cholinergic input instead of all might underestimate the weight of the action (as the local collaterals and inputs from the contralateral PPN were not stimulated), we also showed that cholinergic inputs of the PPN can successfully inhibit the M-current of cholinergic neurons. We concluded that optogenetic activation of a single cholinergic input of the PPN can effectively inhibit the M-current.

Taken together, it seems to be likely that cholinergic activation of a cholinergic nucleus desynchronizes its neuronal population. As the PPN provides local axon collaterals for itself, innervates the contralateral PPN and receives cholinergic fibers from the LDT, cholinergic activation might spread to all mesopontine cholinergic structures and contributes to the desynchronization of cholinergic neuronal populations (Mena-Segovia et al, 2008; Honda and Semba, 1995). Desynchronization of PPN units takes place in parallel with cortical desynchronization (Boucetta et al, 2014; Mena-Segovia et al, 2008; Petzold et al, 2015). One can assume that cholinergic inhibition of the M-current contributes to desynchronization of cholinergic structures, and in turn, to regulation of global brain states.

Neuronal M-current is mostly generated by the combination of KCNQ2 and 3 subunits or homomeric KCNQ2 (Wang et al, 1998; Brown and Passmore, 2009) and by KCNQ5 (Shah et al, 2002; Huang ad Trussell, 2011). Heteromers formed by KCNQ2/KCNQ3 subunits are located in the axon initial segments and nodes of Ranvier, regulating firing (Schwarz et al, 2006; Klinger et al, 2011). KCNQ5 is located in postsynaptic membranes of hippocampal neurons, and contributes to network synchronization (Fidzinski et al, 2015) and in presynaptic locations controlling neurotransmitter release (Huang and Trussell, 2011). KCNQ3/KCNQ5 heteromers also exist in the CNS (Jentsch, 2000; Delmas and Brown, 2005). KCNQ4 in CNS has a more restricted pattern than the other subunits, which adds more variability to the function of the M-current in neurons. We checked the hypothesis whether KCNQ4 subunit contributes to the M-current of the PPN. It was shown that ‒out of its important locations in the periphery (cochlear outer hair cells, DRG, Merkel- and Pacinian corpuscules (Kubisch, 1999; Heidenreich, Lechner et al, 2011) its expression is restricted to brainstem auditory nuclei and the RAS (Kharkovets et al, 2000). From the latter group of nuclei, the VTA and the raphe nuclei had intense labelling, but ‒although not pointed out in the paper-moderate labelling was seen in the PPN, as well. Although the presence of the subunit in the VTA and raphe nuclei was shown with morphological methods, the functional presence of the subunit was only recently confirmed (Su et al, 2019; McGuier et al, 2018; Zhao et al, 2017). We used morphological and pharmacological methods, as well as transgenic animals to demonstrate the presence of the KCNQ4 subunit in the PPN. All methods uniformly showed that not all PPN cholinergic neurons, but a subset of them expressed KCNQ4 in a proportion in between 9 to 62.5%. This finding is similar to the results of Heidenreich et al. (2011) in the DRG, where also only a fraction of neurons was positive to KCNQ4, as well as in the dorsal raphe (Zhao et al, 2017). However, further studies are needed to determine whether KCNQ4-expressing cholinergic neurons have different roles than KCNQ4-negative ones. It is an intriguing finding that not all PPN neurons, but rather, a smaller caudal population possess it. One can hypothesize that the presence of KCNQ4 subunit is characteristic for certain sensory pathways and to nuclei modulating them for adaptation to sleep-wakefulness cycles (Kharkovets et al, 2000; Heidenreich et al, 2011).

A noteworthy discrepancy was seen between the proportion of neurons possessing KCNQ4 subunits (9%) and the number of neurons having functional KCNQ4 subunits revealed by subunit-specific openers (14-27%), as well as the proportion of neurons lacking M-current in KCNQ4 KO mice (62.5%). The great differences between the proportions are probably due to the alteration in expression of other KCNQ subunits in KCNQ4 KO mice. Besides the lack of KCNQ4 mRNA, KCNQ5 mRNA was completely missing and a reduction of KCNQ2 together with a relative increase in KCNQ3 was seen. Alterations in the expression pattern of KCNQ channel subunits were observed in different tissues during pathologies such as hypertension, vascular tumors or retinal degeneration, which in turn contributes to alterations in cellular properties (Jepps et al, 2011; Caminos et al, 2015; Serrano-Novillo et al, 2020). In conclusion, these observations rise the possibility that KCNQ4 is not only important as one of the subunits forming channels for M-current, but might be a potential regulator of the expression of other M-current subunits, setting its physiological function. In agreement with this, RAS-related behavioral alterations were obtained in KCNQ4 KO mice (see below).

In this project, we found that deletion of KCNQ4 subunit, partially due to its expression in the PPN, caused disturbances in adaptation to alterations in the circadian rhythm. In complete darkness, KCNQ4 KO mice increased the activity time, accompanied with longer distances ran under this condition compared to WT littermates. In consequence, KCNQ4 KO mice showed a lower capacity to regulate sleep-wakefulness cycles upon changes in environmental light conditions, suggesting a role in regulating RAS function. In 2-months-old animals, the hearing loss is already present but less prominent than in older ones (Carignano et al, 2019), thus, actions seen on activity cycles are potentially at least partially due to the lack of the ion channel subunit in brainstem structures.

Another argument for this is that there is no significant difference in activity cycles and movement in alternating light-darkness conditions. Changes in outer light conditions possibly do not affect hearing and tactile sensation, rather the structures regulating activity cycles. It is also known that raphe nuclei and the VTA have functional KCNQ4 subunits, which affect neuronal functions in this region (Su et al, 2019; McGuier et al, 2018; Zhao et al, 2017). One can hypothesize that these structures -as they are members of the RAS-contribute to the alterations in activity cycles found in KCNQ4 KO animals. The PPN, the raphe nuclei and the VTA are all involved in both, sleep-wakefulness and movement regulation (Jing et al, 2019; Su et al, 2019; McGuier et al, 2018; Zhao et al, 2017; Monti, 2010; MacLaren et al, 2018; Mena-Segovia and Bolam, 2017). Therefore, as it was expected, distances run under different environmental light conditions were significantly different in WT and KO animals. Based on our findings and literature data, we can conclude that potassium channels containing KCNQ4 subunit in the RAS including the PPN are modulating activity cycles and movement. Symptoms related to this are potentially present on DFNA2 patients, but ‒to the best of our knowledge-it has been not described yet. Besides, KCNQ4 expression in human CNS has not been investigated so far.

The main pathology generated by KCNQ4 malfunction is the progressive hearing loss DFNA2. Although it is considered as non-syndromic, Heidenreich et al. (2011) demonstrated another symptom. They showed that tactile sensation is enhanced both, in animal models and in DFNA2 patients. Our results suggest that ‒besides the dominating hearing loss and the additional changes of tactile sensation-there are further potential symptoms of DFNA2.

One can conclude that the KCNQ4-dependent M-current of the RAS might have a modulatory role on adaptation of activity cycles to environmental changes and on the related changes in the motor activity. The M-current of the PPN has an important contribution to it, as it is known about the nucleus that it regulates sleep-wakefulness cycles and motor activity. Although the KCNQ4 subunit is not the only KCNQ subunit of the nucleus, it has a significant contribution to functions of the M-current. As KCNQ4 is selectively expressed in certain brainstem nuclei, could be a pharmacological target for selective modulation of sleep-wakefulness cycles. KCNQ4-specific openers were hypothesized as effective drugs for treatment of psychiatric diseases (Sotty et al, 2009; Zhao et al, 2017; Su et al, 2019; McGuier et al, 2018). In addition, based on its actions on the PPN, these openers might affect sleep-wakefulness cycles; as well as having the potential for slowing progression of neurodegenerative diseases affecting cholinergic neurons of the PPN (MacLaren et al, 2018) or helping adaptation to artificial alterations of the circadian rhythm (e.g. jet lag or space travel; Brainard et al, 2016). Furthermore, one can also hypothesize that DFNA2 is possibly not a non-syndromic hearing loss, as alterations in sleep-wakefulness cycles and movement regulation might be seen. These latter hypotheses need further confirmation by clinical studies.

## Materials and methods

### Solutions, chemicals

For electrophysiological experiments, artificial cerebrospinal fluid (aCSF) was used in the composition below (in mM): NaCl, 120; KCl, 2.5; NaHCO_3_, 26; glucose, 10; myo-inositol, 3; NaH_2_PO_4_, 1.25; sodium-pyruvate, 2; CaCl2, 2; MgCl_2_, 1; ascorbic acid, 0.5; pH 7.4. For preparation of slices, low Na^+^ aCSF was administered. In this solution, 95 mM NaCl was replaced by sucrose (130 mM) and glycerol (60 mM). All chemicals were purchased from Sigma (St. Louis, MO, USA), unless stated otherwise.

### Animals, preparation

Animal experiments were conducted in accordance with the appropriate national and international laws (EU Directive 2010/63/EU for animal experiments) and institutional guidelines on the care of research animals. The experimental protocols used below were approved by the Committee of Animal Research of the University of Debrecen (6/2011/DEMÁB; 5/2015/DEMÁB; 3-1/2019/DEMÁB) and Universidad Nacional del Sur (083/2016). 10-19 days old mice were employed for slice electrophysiology, whereas 52-69 days old mice were used for behavioral tests. Mice expressing tdTomato fluorescent protein or channelrhodopsin2 (ChR2) in a choline acetyltransferase-(ChAT) dependent way (n = 59 and 5, respectively), as well as mice expressing tdTomato in a type 2 vesicular glutamate transporter (Vglut2) dependent way (n = 13) from both sexes were employed. In order to obtain mice for slice electrophysiology, homozygous floxed-stop-tdTomato (B6;129S6-Gt(ROSA)26Sor^tm9(CAG-tdTomato)Hze/^J; Jax mice accession number: 007905) and ChAT-cre (B6;129S6-Chat^tm2(cre)Lowl/^J; Jax number: 006410) and Vglut2-cre (*Slc17a6*^*tm2(cre)Lowl*^ (also called Vglut2-ires-Cre); Jax number: 028863); or homozygous floxed-stop-channelrhodopsin-2 (B6;129S-Gt(ROSA)26Sortm32.1(CAG-COP4*H134R/EYFP)Hze/J) and ChAT-cre (see above) strains purchased from Jackson Laboratories (Bar Harbor, ME, USA) were crossed in our animal facility. The KCNQ4-knockout (KO) strain was obtained from Prof. Thomas Jentsch (Kharkovets et al, 2006). Heterozygous animals were bred in the animal facility of the Department of Physiology, University of Debrecen. Young pups were genotyped, knockout animals and wild type littermates were employed. In total, 10 KCNQ4 knockout and 12 wild type mice were used for slice electrophysiology and 13 knockout and 15 wild type mice was employed for behavioral tests.

200 μm thick coronal midbrain slices were prepared in ice-cold (cca. 0 - −2 °C) low Na^+^ aCSF with a Microm HM 650V vibratome (Microm International GmbH, Walldorf, Germany). The slices were incubated in normal aCSF on 37°C for 1 hour before starting recording.

### 2.3. Electrophysiology

Patch pipettes with resistance of 6-8 MΩ were fabricated, and filled with internal solution with the following composition (in mM): K-gluconate, 120; NaCl, 5; 4-(2-hydroxyethyl)-1-piperazineethanesulfonic acid (HEPES), 10; Na_2_-phosphocreatinine, 10; EGTA, 2; CaCl_2_, 0.1; Mg-ATP, 5; Na_3_-GTP, 0.3; biocytin, 8; pH 7.3. Whole-cell patch-clamp experiments were conducted at room temperature (22-25°C) on neuronal somata (and in some cases, on proximal dendrites 20-35 μm distal from the soma) with an Axopatch 200A amplifier (Molecular Devices, Union City, CA, USA). Clampex 10.0 software (Molecular Devices, Union City, CA, USA) was used for data acquisition and Clampfit 10.0 (Molecular Devices) software for data analysis. Only stable recordings with minimal leak currents were considered and recordings with series resistance below 30 MΩ, with less than 10% change were included.

Both voltage- and current clamp configurations were used. In certain experiments, 1 μM tetrodotoxin (TTX; Alomone Laboratories, Jerusalem, Israel) was used to eliminate action potential generation. For blockade of M-current, 20 μM XE991 (10,10-*bis*(4-Pyridinylmethyl)-9(10*H*)-anthracenone dihydrochloride; Tocris Cookson Ltd., Bristol, UK) was used. M-current openers, as the non-specific retigabine, the KCNQ2- and KCNQ4-specific ML213 (*N*-(2,4,6-Trimethylphenyl)-bicyclo[2.2.1]heptane-2-carboxamide) and the KCNQ2- and KCNQ3-specific ICA27243 (N-(6-chloro-pyridin-3-yl)-3,4-difluoro-benzamide; 20 μM; Tocris Cookson Ltd., Bristol, UK) were administered in certain experiments (Gunthorpe et al, 2012; Linley et al, 2012; Brueggemann et al, 2014; Wickenden et al, 2008).

Protocols detailed below were used to assess the presence of M-current or the functional consequences of the M-current on PPN cholinergic (and, in some cases, glutamatergic) neurons. For determining spike train properties, current clamp configuration was used. For recording of the M-current, voltage clamp configuration was used. Neurons were held on −20 mV holding potential and 1-s-long repolarizing steps were employed from −30 to −60 mV with 10 mV decrement. Recordings were performed with TTX, except when the laterodorsal tegmental nucleus was stimulated with optogenetic methods. For detection of spike train pattern, 1-s-long square current pulses were used between −30 pA and +120 pA with 10 pA increment in current clamp configuration. The resting membrane potential was set to −60 mV. Adaptation index (AI) was calculated by using the following formula: AI = 1−(F_last_/F_initial_), where F_last_ is the frequency of the last 2 action potentials and F_initial_ is the average frequency of the first 3 action potentials. Only those recordings were considered where at least 8 action potentials were seen in control. In those cases where this number decreased below 5 due to the robust action of M-current openers, AI was considered as 1.

When synchronization in action potential firing of neighboring neurons was assessed, absolute values of time differences between action potentials were plotted with using 20 ms bins. The individual graphs under control conditions and after application of XE991 were fit with single exponential function (*y = a + b*e*^*(τ*x)*^), and the parameter ‘τ’ was used as a measure of synchronicity.

In certain cases, simultaneous recordings were performed on neuronal somata and proximal dendrites (20-35 μm away from the soma) using patch pipettes with 6-8 MΩ resistance. In other cases, neighboring but synaptically non-coupled cholinergic neurons were patched and both neurons were simultaneously depolarized by using 1-s-long depolarizing square current injections with an amplitude of 100 pA in current clamp configuration.

In experiments with optogenetics, 500 μm thick coronal midbrain slices were cut including the laterodorsal tegmental nucleus (LDT) and the pedunculopontine nucleus (PPN). An optical fiber was administered to the LDT and 50 ms long illuminations with 470 nm wavelength and 1 Hz frequency were used. Before and in parallel with it, whole cell patch clamp recordings of the M-current (see above) were performed on PPN cholinergic neurons.

Visualization of the tdTomato and EYFP fluorescent markers was performed by using a widefield fluorescent imaging system (Till Photonics GmbH, Gräfeling, Germany) containing a xenon bulb-based Polychrome V light source, a CCD camera (SensiCam, PCO AG, Kelheim, Germany), an imaging control unit, and the Till Vision software (version 4.0.1.3).

### 2.4. Morphological identification of the investigated neurons

Neurons were labeled with biocytin during patch clamp recording and slices were fixed (4% paraformaldehyde in 0.1 M phosphate buffer; *pH* 7.4; 4 °C) for morphological analysis of the neurons. Tris buffered saline (in mM, Tris base, 8; Trisma HCl, 42; NaCl, 150; *pH* 7.4) supplemented with 0.1% Triton X-100 and 10% bovine serum (60 min) was applied for permeabilization. For recovery, samples were incubated in streptavidin-conjugated Alexa488 (1:300; Molecular Probes Inc., Eugene, OR, USA) dissolved in phosphate buffer for 90 min.

After the recovery procedure, neurons were visualized with confocal microscope (Zeiss LSM 510; Carl Zeiss AG, Oberkochen, Germany); tile scan images were taken with 40x objective and with 1 μm optical slices.

### 2.5. Immunohistochemistry

KCNQ4 and choline acetyltransferase (ChAT) immunofluorescence was performed on 15 μm coronal brain sections from cryostat of adult (3-6 months old) transcardially perfused WT, KCNQ4-KO, ChAT-tdTomato or Vglut2-tdTomato mice. Brain samples were post-fixed in 4% paraformaldehyde for 24 hours. After 3×10 minutes incubation with PBS, the slices were permeabilized with 2% Nonidet and 1% BSA for 1 hour. Anti-choline acetyltransferase (anti-ChAT) primary antibody from goat (Millipore, Temecula, CA, USA) was incubated for 48 hours in 1:100, with anti-KCNQ4 1:400 from rabbit. Donkey anti-goat Alexa 555 and Donkey anti-rabbit Alexa 488 antibodies (Vector Laboratories Inc., Burlingame, CA, USA) were used for another 24 h. After completion of the protocol above, confocal images were taken in a similar way described in Chapter 2.4.

### 2.6. RNA extraction, Reverse Transcription and qPCR

Coronal midbrain blocks were prepared and areas containing the PPN were taken out from adult (15-30 weeks old) mice. For each experiment, samples from three to four mice were pooled. Total RNA was extracted from PPN was extracted using the TransZol reagent (TransGen Biotech, China) in combination with the Direct-Zol RNA mini prep kit (Zymo Research, USA). cDNA was produced from 500 ng of total RNA with EasyScript Reverse Transcriptase (TransGen Biotech Cat #AE101) using anchored oligo (dT)s following manufacturer’s indications. Quantitative PCR (qPCR) was carried out using the cDNA generated previously employing the SensiFAST SYBR mix No-Rox Kit (Bioline, UK in a Rotor-Gene 6000 real-time PCR cycler (Corbett Research, USA). The primer used were listed in Table 2. HPRT was used as reference gene. Data analysis was done applying the ΔCt and ΔΔCt method (Livak and Schmittgen, 2001; Schmittgen and Livak, 2008) to obtain relative mRNA expression. All data was represented as mean ± SD. One-way ANOVA and post hoc Tukey test were applied for comparison of the results obtained from three independent experiments done in triplicate.

**Table 2.**
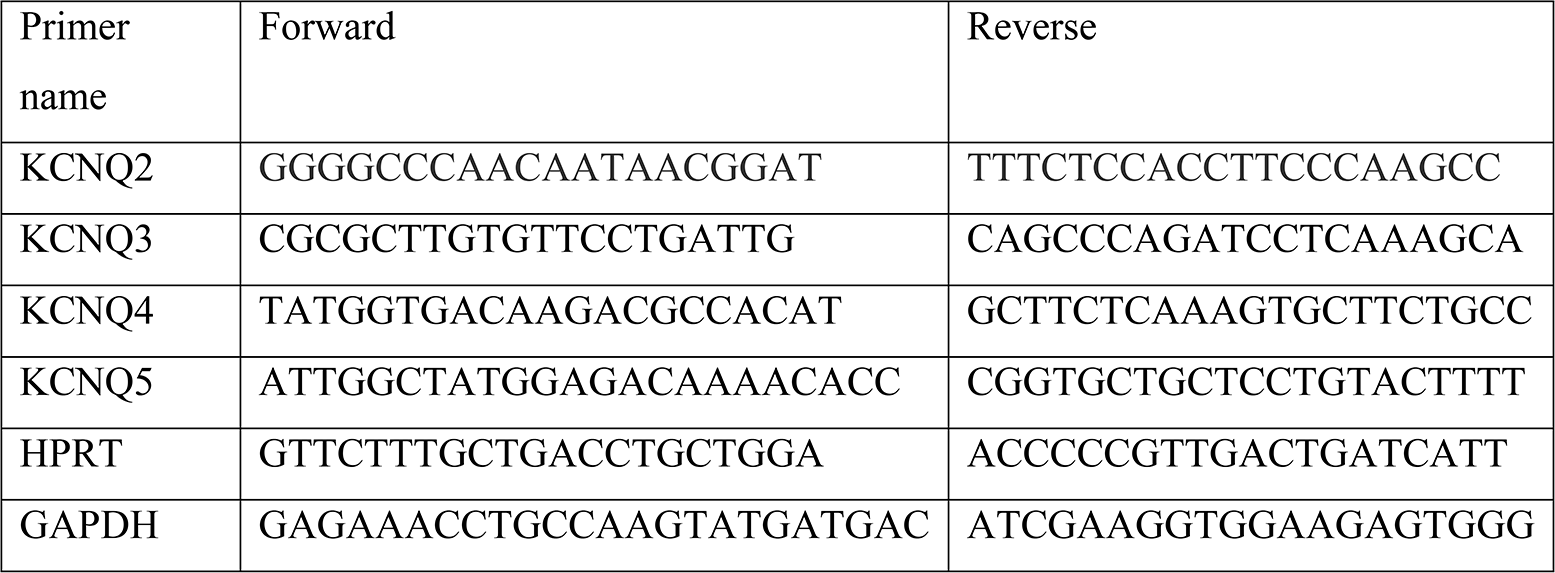
Primers used for qPCR experiments.

### 2.7. Activity wheel test

Duration of activity cycles and distances moved per day were checked by activity wheel test (Campden Instruments Ltd., Loughborough, UK). Young adult (52-69 days old) KCNQ4 knockout mice (n = 16) and wild type littermates (n =16) were used. Mice were housed individually in cages with activity wheel. As it was previously known that there is a progress in deafness of KCNQ4 KO mice and the activity wheel performance also changes with age, we aimed to use as young animals as possible and to minimize the progress during the experiment by setting the experimental animals to shorter circadian periods (Kharkovets et al, 2006; Fodor et al, 2020). Animals had voluntary and unlimited access to the activity wheel and placed in a room for 7 days to accommodate to alternating 6 h illumination and 6 h darkness periods. After accommodation, 5 days of recording was done with the same conditions of illumination (LD conditions). After that, recording was continued during 5 days of complete darkness (DD conditions). The last 3 days of recordings were taken into account. Period time was determined by Lomb-Scargle periodogram (www.circadian.org). Due to technical problems, 3 of the knockout and 1 of the wild type animals were taken out from the study, thus 13 knockout and 15 wild type mice were involved.

### 2.8. Statistics

All data represent mean ± SEM. The normal distribution of the datasets was evaluated with D’Agostino and Pearson omnibus normality test. Paired Student’s *t*-test, one way ANOVA and post hoc Tukey Multiple Comparison test were applied for assessing statistical significance for pairwise comparison in case if the datasets had normal distribution, whereas Bonferroni’s multiple comparison test was employed for multiple comparisons. Statistical analysis was performed using Microsoft Excel or SigmaPlot 12.0 and GraphPad Prism 5.01 (GraphPad software, San Diego, USA). The level of significance was p < 0.05.

## Acknowledgements

This work was supported by the Hungarian National Brain Research Program (KTIA_13_NAP-A-I/10.; BP); the Argentinian-Hungarian Scientific and Technological Cooperation Grant (TÉT_15-1-2016-0087; GS and BP) and the Thematic Excellence Programme of the Ministry for Innovation and Technology in Hungary (Space Sciences tematical programme of the University of Debrecen; ED_18-1-2019-0028). TB was supported by the Stipendium Hungaricum PhD programme. The KCNQ4 knockout strain was a kind gift from Prof. Thomas Jentsch (FMP/MDC, Berlin, Germany). The authors are indebted to Dr. Rita Gálosi (University of Pécs, Hungary) for her valuable comments.

## Author contribution

TB, KA, CsA, and PK performed patch clamp and optogenetic experiments and contributed to genotyping and maintenance of transgenic strains. SS and LD performed qPCR and immunohistochemical (IHC) experiments. SzP and BP performed the activity wheel test. TB, SS, LD, GS and BP wrote the manuscript. BP supervised the patch clamp and behavioral studies, GS supervised the qPCR and IHC studies.

## Conflict of interest

The authors declare no conflict of interest.

